# Visualizing the pH in *Escherichia coli* colonies *via* the sensor protein mCherryEA allows high-throughput screening of mutant libraries

**DOI:** 10.1101/2021.07.08.451719

**Authors:** Fabian Stefan Franz Hartmann, Tamara Weiss, Jing Shen, Dóra Smahajcsik, Gerd Michael Seibold

## Abstract

Cytoplasmic pH is tightly regulated by diverse active mechanisms and interconnected regulatory processes in bacteria. Many processes and regulators underlying pH-homeostasis have been identified *via* phenotypic screening of strain libraries towards non-growth at low or high pH values. Direct screens with respect to changes of the internal pH in mutant strain collections are limited by laborious methods including fluorescent dyes or radioactive probes. Genetically encoded biosensors equip single organisms or strain libraries with an internal sensor molecule already during the generation of the strain. In this study, we used the pH-sensitive mCherry variant mCherryEA as ratiometric pH biosensor. We visualized the internal pH of *E. coli* colonies on agar plates by the use of a Gel-Doc imaging system. Combining this imaging technology with robot-assisted colony picking and spotting allowed us to screen and select mutants with altered internal pH values from a small transposon mutagenesis derived *E. coli* library. Identification of the TN- insertion sites in strains with altered internal pH levels revealed that the transposon was inserted into *trkH* (encoding a transmembrane protein of the potassium uptake system) or the *rssB* gene (encoding the anti-adaptor protein RssB which mediates the proteolytic degradation of the general stress response regulator RpoS), two genes known to be associated with pH-homeostasis and pH stress adaptation. This successful screening approach demonstrates that the pH- sensor based analysis of arrayed colonies on agar plates is a sensitive approach for the fast identification of genes involved in pH-homeostasis or pH stress adaptation in *E. coli*.

**Importance:** Phenotypic screening of strain libraries on agar plates has become a versatile tool to understand gene functions and to optimize biotechnological platform organisms. Screening is supported by genetically encoded biosensors that allow to easily measure intracellular processes. For this purpose, transcription-factor-based biosensors have emerged as the sensor-type of choice. Here, the target stimulus initiates the activation of a response gene (e.g. a fluorescent protein) followed by transcription, translation and maturation. Due to this mechanistic principle, biosensor readouts are delayed and cannot report the actual intracellular state of the cell in real-time. To capture fast intracellular processes adequately, fluorescent reporter proteins are extensively applied. But these sensor-types are not utilized for phenotypic screenings so far. To take advantage of their properties, we here established an imaging method, which allows to apply a fast ratiometric sensor protein for assessing the internal pH of colonies in a high-thoughput manner.

## 1 Introduction

Genetically encoded sensors targeting intracellular metabolites have become a versatile tool for physiological studies in diverse organisms (Sanford and Palmer, 2017; Koch *et al.*, 2019; Shin *et al.*, 2020). These sensors have been successfully applied in bacteria for screening optimized production strains, activity of/or resistance against antimicrobial compounds as well as for assessing physiological states and metabolic fluxes (Schallmey *et al.*, 2014; Maglica *et al.*, 2015; Crauwels *et al.*, 2018; Monteiro *et al.*, 2019; Heins *et al.*, 2020). Commonly, two different types of genetically encoded sensors are used: transcription-factor-based biosensors (TFBs) or fluorescent-reporter-proteins (FRPs). TFBs are the most extensively developed and applied biosensors due to their simplicity to engineer. The basic design generally relies on transcription factors, which natively react to effectors (activator or repressor). Upon interaction, the TF-effector complex targets or releases a cognate promotor sequence to transduce a response through activation or repression of the respective downstream reporter gene such as a fluorescent protein. Dynamics of TFBs towards monitoring changes of target product concentrations in real-time are limited as the sensor signal depends on transcription, translation, maturation of the fluorescent protein or degradation. This, however, provides the advantage that the sensor signal is stable even when exposing a sensor strain to varying external conditions, making these type of sensors suited for high-throughput screening of strains *via* e.g. fluorescence activated cell sorting (FACS) (Schallmey *et al.*, 2014; Eggeling *et al.*, 2015). In contrast, FRPs respond in real-time to alterations of internal target parameters or metabolites such as pH, ATP or NADPH (Goldbeck *et al.*, 2018; Reyes-Fernández and Schuldiner, 2020; Botman *et al.*, 2020; Deng *et al.*, 2021). Here, an already produced sensor protein undergoes analyte-dependent conformational changes accompanied by a change of the fluorescence properties (Bermejo *et al.*, 2011b; Martynov *et al.*, 2018). Consequently, FRPs were successfully applied for real-time monitoring of internal metabolite levels or oxidation states upon externally applied perturbations (Bermejo *et al.*, 2011b; Martinez *et al.*, 2012; Goldbeck *et al.*, 2018; Martynov *et al.*, 2018; Depaoli *et al.*, 2019; Hartmann *et al.*, 2020). Measuring internal parameters of individual microbial strains of mutant collections *via* FRPs could benefit by the fact that actual values can be measured rather than events which occurred in the past. Furthermore, fast sensor dynamics would allow the scientist to perform a sensor calibration and validation of sensor properties at the level of the actual screening. However, applying FRPs for FACS-based high-throughput screening of microbial strains is challenging due to the varying external conditions. To take advantage of the properties of FRPs, the screening method needs to allow maintaining constant conditions when conducting the sensor analysis, such as phenotypic screening on agar plates.

Many bacterial regulatory mechanisms have been identified *via* phenotypic screening of strain libraries with respect to its growth patterns under different conditions. Screening of *E. coli* and other microorganisms towards growth vs. non-growth at low or high pH-values revealed many of the processes and control mechanisms underlying pH-homeostasis (Reva *et al.*, 2006; Mira *et al.*, 2010; Guerrero-Castro *et al.*, 2018; Palud *et al.*, 2018; Bushell *et al.*, 2021). To achieve pH homeostasis, *E. coli* possesses regulatory networks for acid and alkaline conditions, which trigger expression of distinct sets of genes (Maurer *et al.*, 2005; Hayes *et al.*, 2006). For response to acid conditions, *E. coli* activates systems for consumption of intracellular protons *via* deamination and decarboxylation of amino acids, formation of neutralizing ammonia from glutamine and extrusion of protons *via* the F_1_Fo-ATPase (Lund *et al.*, 2014; Guan and Liu, 2020; Lund *et al.*, 2020). Moreover, potassium uptake and accumulation was shown to be essential for the maintenance of internal pH in *E. coli*. Under acidic conditions a neutral pH in the cytoplasm can only be maintained if sufficient potassium is available, accumulated *via* one of three potassium uptake systems (Roe *et al.*, 2000; Reyes-Fernández and Schuldiner, 2020). Upon exposure to alkaline pH, *E. coli* expresses genes for cation proton antiporters, which import protons in exchange for sodium and/or potassium ions (Krulwich *et al.*, 2011; Ito *et al.*, 2017). Following the identification of a mutant strain possessing a pH-dependent growth phenotype, the cytoplasmic pH of the isolated mutant is measured *via* fluorescent dyes (e.g. BCECF and SNARF), radioactive probes (Kashket, 1985; Han and Burgess, 2010) or genetically encoded sensor proteins (Martynov *et al.*, 2018; Rajendran *et al.*, 2018). For this purpose, different ratiometric pH responsive RFPs such as pHluorin and pHred have been developed, both possessing a pK_a_ of 6.9 but different intrinsic fluorescence properties (Miesenböck *et al.*, 1998; Tantama *et al.*, 2011; Martynov *et al.*, 2018). Recently, the mCherry variant mCherryEA was shown to be an effective ratiometric red fluorescent protein pH biosensor possessing a pK_a_ of 7.3 (Rajendran *et al.*, 2018). This is close to the range of internal pH values reported for *E. coli* (7.4-7.9) (Slonczewski *et al.*, 2009), making this sensor protein well suited for applications in *E. coli*.

In this study, we successfully visualized ratiometric sensor signals from the genetically encoded pH sensor mCherryEA in *E. coli* colonies cultivated on agar plates by using an imaging system equipped with filters for fluorescence detection. Combining this imaging technology with robot-assisted colony picking and spotting allowed us to screen and select mutants with altered internal pH levels from a small transposon mutagenesis derived *E. coli* library. We here show that a sensor analysis with the pH sensor mCherryEA of colonies on agar plates is a sensitive approach for the fast identification and characterization of genes involved in pH-homeostasis or pH stress adaption in *E. coli*. The here established approach can easily be adapted for other strain backgrounds or genetically encoded FRPs targeting another product or internal parameter and thus enables novel studies in microbial systems biology.

## 2 Materials and methods

### Strains, media, and culture conditions

Bacterial strains and plasmids used in this study are listed in Table 1. Cloning as well as biosensor expression for the preparation of crude cell extracts was carried out using *E. coli* DH5 α, cultivated in Lysogeny Broth (LB) medium (Bertani, 2004). *E. coli* MG 1655 and *E. coli* TK 2309 were pre-cultured in LK medium (5 g/L yeast extract; 10 g/L BactoTryptone; 6.4 g/L KCl). For main cultures as well as short time cultivations to assess impact of potassium on pH, K_0.1_, K_30_ and K_120_ media was prepared (Epstein and Kim, 1971). For this purpose K_0_ buffer (8.25 g/L Na_2_HPO_4_ *2H_2_O, 2.8 g/L NaH_2_PO_4_*H_2_O, 1 g/L (NH_4_)_2_SO_4_) was prepared and a final buffer concentration of 0.1 mM KCl or 60 mM KCl was adjusted in order to get K_0.1_ and K_30_ media, respectively. K_120_ media was prepared by using K_120_ buffer instead (8 g/L K_2_HPO_4_; 3.1 g/L KH_2_PO_4_; 1 g/L (NH_4_)_2_SO_4_). All media and buffers were prepared using ultrapure water prepared by using an arium^®^pro ultrapure water purification system (Sartorius, Germany). Prior inoculation, all K_x_ media were supplemented with 0.2 % glucose, 0.4 mM MgSO_4_*7H_2_O, 0.6 μM (NH_4_)_2_SO_4_ x FeSO_4_ *6H_2_O and thiamin- HCl 0.0001% (w/v). For screening purposes, Screening Broth (SB) medium (5 g/L yeast extract; 10 g/L BactoTryptone; 100 mM NaCL; 50 mM KCl, buffered with 50 mM TRIS) was used (pH 7.0). For preparation of agar plates 16 g/L agar was added to the respective media. Strains carrying plasmids and transposons were cultivated in presence of kanamycin (50 μg/mL) or chloramphenicol (20 μg/mL). If required, 1 mM IPTG was added to induce expression of the gene for the biosensor. For fluorescence imaging of agarose plates, 50 mL of the medium were used for each plate (SBS-format PlusPlates, Singer Instruments, United Kingdom) and supplemented with black food dye (30 μL/plate) to reduce the fluorescence background from the media.

**Table 1:**
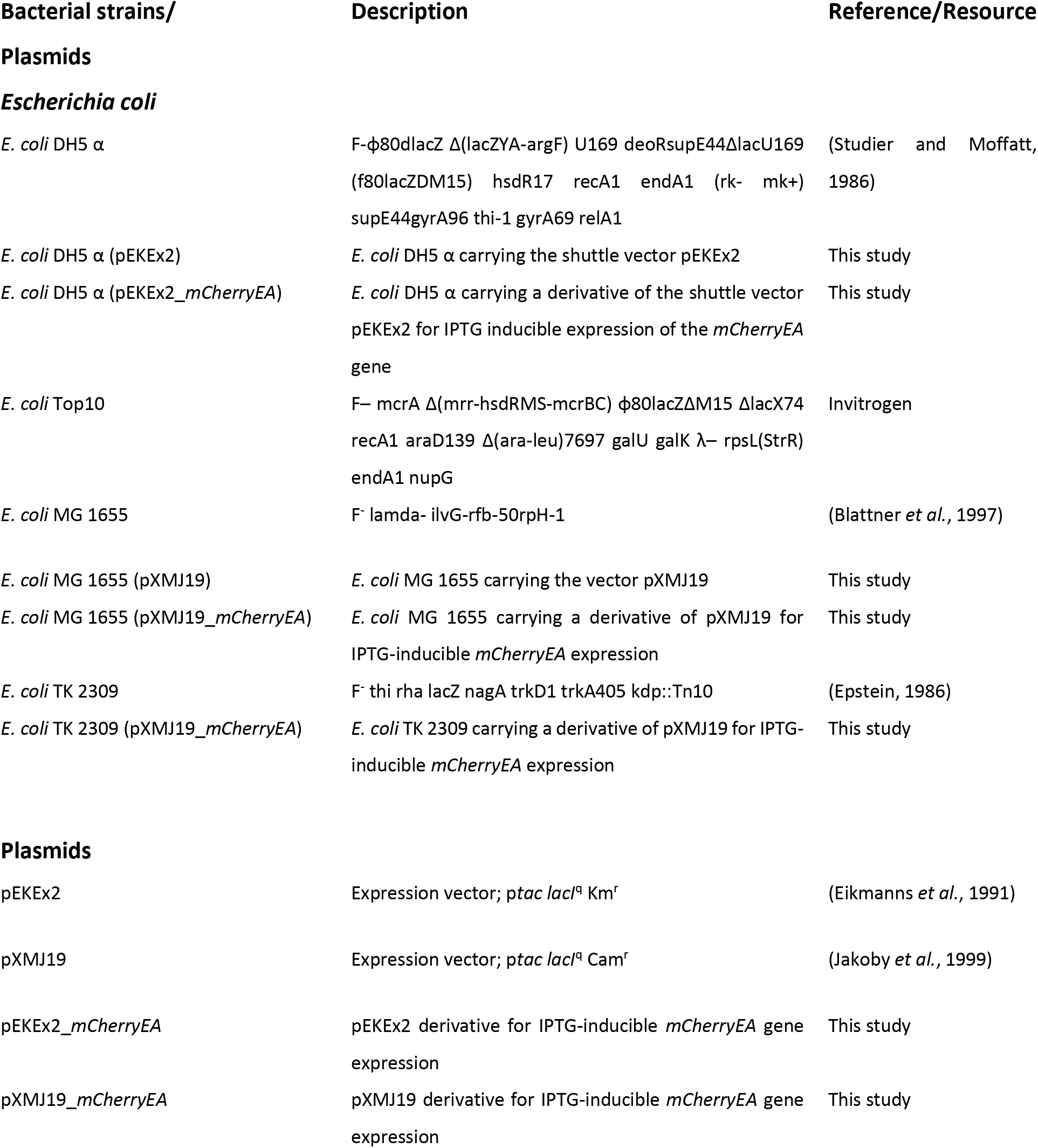
Bacterial strains and plasmids used in this study.

### Genetics

Gene fragment synthesis was carried out by Integrated DNA Technologies (IDT) (Denmark). The sequence is provided in Table S1. For amplification of the gene fragments primer sets were designed using NEBuilder. All primers are listed in Table S1. Expression plasmids were assembled using the Gibson assembly kit (New England Biolabs, U.S.A) according to the manufacturer’s instructions. After transformation, recombinant strains were selected using LB-agar plates supplemented with appropriate antibiotics. Recombinant *E. coli* MG1655 and TK 2309 strains were selected on LK-media agar plates with appropriate antibiotics. The plasmids were analyzed *via* PCR, restriction digests, and DNA sequencing (Eurofins Genomics, Germany).

### Transposon mutagenesis library generation and introduction of pXMJ19-mCherryEA into the library

The EZ-Tn5 <KAN-2> Tnp transposome (Epicentre Biotechnologies, U.S.A) was introduced into *E. coli* Top10 by electroporation. Cells were subsequently spread plated on LB agar plates containing 50 μg/mL kanamycin and incubated overnight at 37 °C. Single colonies from the spread plates were analyzed to determine the diversity of insertion sites *via* linker PCR as described below. Pools of 5-6 × 10^3^ colonies were collected and frozen at −80 °C in 0.9 % NaCl containing 30 % glycerol. For the introduction of the sensor plasmid, 100 μL of these glycerol stocks were used to inoculate 2 mL LB containing 50 μg/mL kanamycin and cultivated overnight on a shaker at 180 rpm and 37 °C. Then, 2 mL of the pre-culture were used to inoculate 50 mL LB containing 50 μg/mL kanamycin in a 500 mL flask, cultivated at 200 rpm, 37 °C, until the culture reached an OD_600_ of 0.4, and then competent cells were performed as described in (Sambrook *et al.*, 1989). The plasmid pXMJ19_*mCherryEA* was transformed into the mutant library competent cells using electroporation. Transformants were spread plated on LB plates containing 50 μg/mL kanamycin and 20 μg/mL chloramphenicol and used for further screening.

### Identification of Tn5 insertion sites using Linker PCR

Genomic DNA was extracted from 1.5 mL cultures by using the GenElute™ Bacterial Genomic DNA Kit (Merck, Germany). Linker PCR was used to test individual transformant colonies and to determine the diversity of insertion sites. Genomic DNA (2 μg) was digested with the *AluI* restriction enzyme (New England Biolabs, U.S.A) and purified by using an illustra GFX PCR DNA and Gel Band Purification Kit (GE Healthcare, U.S.A). The linker was generated by annealing 100 μM of each oligonucleotides P2-FW (Table S1) and P3-RV (Table S1) in an annealing buffer (10 mM Tris, 50 mM NaCl, 1 mM EDTA, pH 8.0) after incubation at 95 °C for 2 minutes followed by cooling to 25 °C for 1 hour and chilling to 4 °C. Then the linker was ligated to the ends of restriction fragments (50 ng) by using T4 DNA ligase (New England Biolabs, U.S.A). The ligated DNA templates were cleaned by an illustra GFX PCR DNA and Gel Band Purification Kit (GE Healthcare, U.S.A). Finally, Linker PCR was carried out with the ligated DNA template and transposon-specific oligonucleotides P4-FW (Table S1) and P5-RV (Table S1) by using Phusion X7 and thermocycling conditions of 98 °C for 30 seconds, followed by 35 cycles of 98 °C for 10 sec, 55 °C for 30 seconds and 72 °C for 1 min, with a final extension step of 72 °C for 10 minutes. The resulting PCR samples were run on 2% agarose gels at 100 V for 25 min.

### Fluorescence analysis

Fluorescence measurements of liquid cultures were conducted in black flat-bottomed 96-well microplates (Thermo Fisher Scientific, Germany) using a SpectraMax iD3 multi-mode plate reader (Molecular Devices LLC, U.S.A). Excitation scans were recorded by setting the excitation wavelength between 410 nm and 588 nm and the emission wavelength at 630 nm. For ratiometric analysis of the biosensor signal, the emission maxima obtained upon an excitation at 454 nm and 580 nm were taken and the corresponding biosensor ratio was calculated by dividing the former emission intensity by the latter. Fluorescence imaging was carried out using the photo-documentation system Fusion FX (Vilber Lourmat, France). The Fusion FX was equipped with capsules for excitation at 440 nm and 530 nm with a set exposure time of 40 msec and 1560 msec, respectively. Fluorescence was measured using a 595 nm emission filter. Images were analyzed using the Fusion FX software Evolution-edge provided by Vilber Lourmat.

### *In vitro* and *in situ* characterization of biosensor protein

For *in vitro* characterization, biosensor strains and empty vector controls were cultivated in shaking flasks (50 mL, LB medium with respective antibiotics) until an OD_600_ of 1 was reached. Subsequently, 1 mM IPTG was added in order to induce expression of the gene from the biosensor and cultivation was continued for 16 hours at 37 °C and 180 rpm. For preparation of crude cell extracts, cells were harvested by centrifugation (4000 rpm, 10 min., 4 °C), washed twice in 1 M potassium phosphate buffer with different pH values set by titrating 1 M K_2_HPO_4_ with 1 M KH_2_PO_4_ and finally the washed cells were resuspended in 1 mL of the 1 M potassium phosphate buffer with the respective pH. Disruption of the cells was conducted using a Ribolyzer (Precellys TM Control Device, Bertin Technologies, France) at 6000 rpm, four times for 30 seconds each. Cell debris were removed *via* centrifugation (13.000 g, 20 min.; 4 °C) and 200 μL of the supernatant was transferred to black flat-bottomed 96-well microplates (Thermo Fisher Scientific, Germany) for further fluorescence analysis a SpectraMax iD3 microplate reader was used as stated above.

For *in situ* characterization of the pH biosensor mCherryEA, cells pre-cultivated as described for the *in vitro* characterization were washed with 1 M potassium phosphate buffer (PBS) with different set pH-values and finally resuspended in a PBS with respective pH to an OD_600_ of three. Aliquots of the cell suspensions (190 μL) were then transferred to black flat-bottomed 96-well microplates. Subsequently 10 μL CTAB (0.05 % (w/v)) were added to the wells and the plate was incubated for 5 min. at room temperature in the dark for permeabilization of the cell membrane as previously described (Crauwels *et al.*, 2018; Goldbeck *et al.*, 2018). Subsequently, fluorescence measurements for biosensor analysis were performed in a microplate reader as described above.

### *In vivo* assay to assess pH homeostasis by the use of the plasmid encoded sensor protein mCherryEA in *E. coli* liquid cultures

Single colonies of *E. coli* strains carrying the plasmid pXMJ19_*mCherryEA* were used to inoculate 5 mL LK medium, incubated ON/16 hours and then used to inoculate 50 mL LK medium supplemented with 1 mM IPTG to induce expression of the gene for the pH-biosensor protein mCherryEA. Stationary cells were harvested *via* centrifugation (3500 rpm, 5 min., 4 °C) and washed twice in 0.9 % NaCl. Finally, cells were re-suspended in K_30_ and the suspension then used to inoculate 50 mL K_30_ medium supplemented with 1 mM IPTG. The next day, cells were harvested *via* centrifugation (3500 rpm; 5 min., 4°C) and washed three times with 0.9 % NaCl. Subsequently, cells were grown for 3 hours either in 50 mL K_0.1_ or K_120_-medium supplemented with 1 mM IPTG. Finally, an OD_600_ of 3 was adjusted by re-suspending the cells in fresh K_0.1_ or K_120_ medium (0.2 % glucose (w/v)) with different pH values. Then, 180 μL of each cell suspension was transferred to black 96-well plates. Incubation and fluorescence measurements were carried out using the SpectraMax iD3 plate reader (incubation temperature 37 °C, continuous orbital shaking at medium intensity). Biosensor signals were then recorded in intervals of five minutes for one hour. At the end of the experiment, CTAB (final concentration 0.05 % (v/v)) was added manually to each well in order to verify the biosensor functionality *via* equilibration of external and internal conditions. Moreover, this signal was used for re-calibration of the pH biosensor signals at the end of every experiment.

### Robotic colony picking and spotting

Prior colony picking, wells of 96-well microtiter plates (Greiner bio-one B.V., Netherlands) were filled with 150 μL liquid SB medium. Cells from single colonies on transformation plates were picked up using the colony picking robot (QPix 420, Molecular Devices, LLC, U.S.A.) mounted with a bacterial 96-pin picking head and transferred to the designated well in the 96-well plate. The QPix robot was programmed to create a copy of the target 96-well plate in a second prefilled 96-well plate in order to finally get one working plate and one back-up plate for long-term storage purposes. Following the picking procedure, the plates were incubated for 16 hours in plate holders at 37°C and 280 rpm. Subsequently, 150 μL glycerol (50 % (v/v)) were added to the cultures of the back-up plate, which was then immediately stored at −80°C. The working plate was used as a source plate for robotic spotting using a ROTOR HDA benchtop robot (Singer Instruments, United Kingdom) on rectangular OmnyTray plates (Singer Instruments) with solidified SB medium as the target plate. Four liquid source plates were finally combined on one target solid plate in a 96 well to 384 spots mode. The OmnyTray plates were prepared by using 50 mL of the respective media supplemented with 1 mM IPTG for biosensor expression. Prior liquid to solid spotting, pins were rotated five times in the source 96-well plate in order to generate a homogenous mixture of the cell suspension. Spotting was conducted by setting an overshoot of 1.5 mm and a pin-pressure of 7 % using long 96-well pins (Singer Instruments). To avoid reflection from the plastic edges of the OmnyTray plates as well as effects resulting from the outer barrier of arrayed colonies, the outermost lines and rows of spotted colonies on each plate were excluded from further analysis resulting in 16 x 8 colonies on each target agar plate.

### Data analysis

Analysis of one-way variance (ANOVA) with Tukey’s test was used to assess differences of sensor signals derived from *E. coli* strains harboring the genetically encoded biosensor mCherryEA. Differences were considered statistically significant when *p* < 0.05.

## 3 Results and Discussion

### 3.1 The plasmid encoded sensor protein mCherryEA allows real-time monitoring of internal pH in *E. coli*

The mCherry variant with the I158E and Q160A amino acid exchanges, originally engineered to support excited-state proton transfer for generating a long Stokes shift variant, exhibits at neutral pH two excitation peaks corresponding to the protonated and deprotonated chromophore with a single emission peak (Piatkevich *et al.*, 2010). Based on this property, the mCherry variant named mCherryEA, was found to function as a ratiometric pH sensor protein, because the protonation state of Glu158 is sensitive to the pH of the surrounding solution, which results in pH-dependent protonation of the chromophore (Rajendran *et al.*, 2018). To generate a sensor plasmid encoding the mCherryEA, the gene was synthesized and cloned into the backbone of the expression plasmid pEKEx2, resulting in the plasmid pEKEx2_mCherryEA. Analysis of the fluorescence properties of the pH biosensor protein mCherryEA in cell free extracts of *E. coli* DH5α (pEKEx2_mCherryEA) at different pH-values has shown the expected ratiometric and pH-dependent change of the emission intensity at 630 nm (Fig. 1a), as previously described (Rajendran *et al.*, 2018). In detail, an increase of the pH was accompanied by an increased emission intensity obtained for an excitation at 454 nm (maximum), whereas the emission intensity upon excitation at 580 nm decreased (Fig. 1a). Consistently, no pH-dependent changes of fluorescence were detected in cell-free extracts of the empty vector control strain *E. coli* (pEKEx2) (Fig. S1). Based on the changes of fluorescence emissions for an excitation of both 454 nm as well as 580 nm, the pH dependent ratiometric response of the biosensor mCherryEA was calculated (Fig. 1b). As depicted in Fig. 1b, an approximately eight-fold increase of the ratio occurred with increasing pH values from 6.5 to 9.0. Taking into consideration that internal pH values between 7.4-7.9 have been reported for *E. coli* (Roe *et al.*, 2000; Slonczewski *et al.*, 2009; Krulwich *et al.*, 2011), the mCherryEA biosensor properties seem well suited to assess changes towards both more alkaline as well as acidic internal pH values. For also testing the *in vivo* functionality of the mCherryEA biosensor, *E. coli* DH5α (pEKEx2_mCherryEA) cells were suspended in PBS buffer with different pH values and subsequently the ratios of the fluorescence signals for the biosensor mCherryEA determined for each of the cell suspensions in the microplate reader. As depicted in Fig 1c, increased ratios were determined for *E. coli* DH5α (pEKEx2_mCherryEA) suspensions in high pH PBS buffer and low ratios for suspensions in low pH-values. The ratiometric biosensor curve obtained from mCherryEA in cell-free extracts (Fig. 1b) differed from these determined for suspensions of intact cells (Fig. 1c). This observation indicates that pH-homeostasis might proceed in the intact cells, causing the internal pH to be different from the external. The addition of the quaternary ammonium surfactant cetyltrimethylammonium bromide (CTAB) to cells permeabilizes the cell membrane and disrupts the proton gradient across the cytoplasmic membrane allowing the internal pH to become identical to the external (Cella *et al.*, 1952; Crauwels *et al.*, 2018). Indeed, the addition of CTAB to the suspensions of *E. coli* DH5α (pEKEx2_mCherryEA) with different pH-values resulted in a shift of the ratiometric mCherryEA biosensor signals (Fig. 1c). When the pH_in_ values for the CTAB treated cell suspensions were calculated based on the obtained ratios (Fig. 1d), pH_in_ values within the expected dynamic range of the biosensor were obtained (Fig. 1d). The pH_in_ for non-permeabilized cell suspensions of *E. coli* DH5α (pEKEx2_mCherryEA), however, was different from the external pH (Fig. 1d), indicating that the cells probably performed to some extent pH-homeostasis even in absence of an energy source.

**Figure 1:**
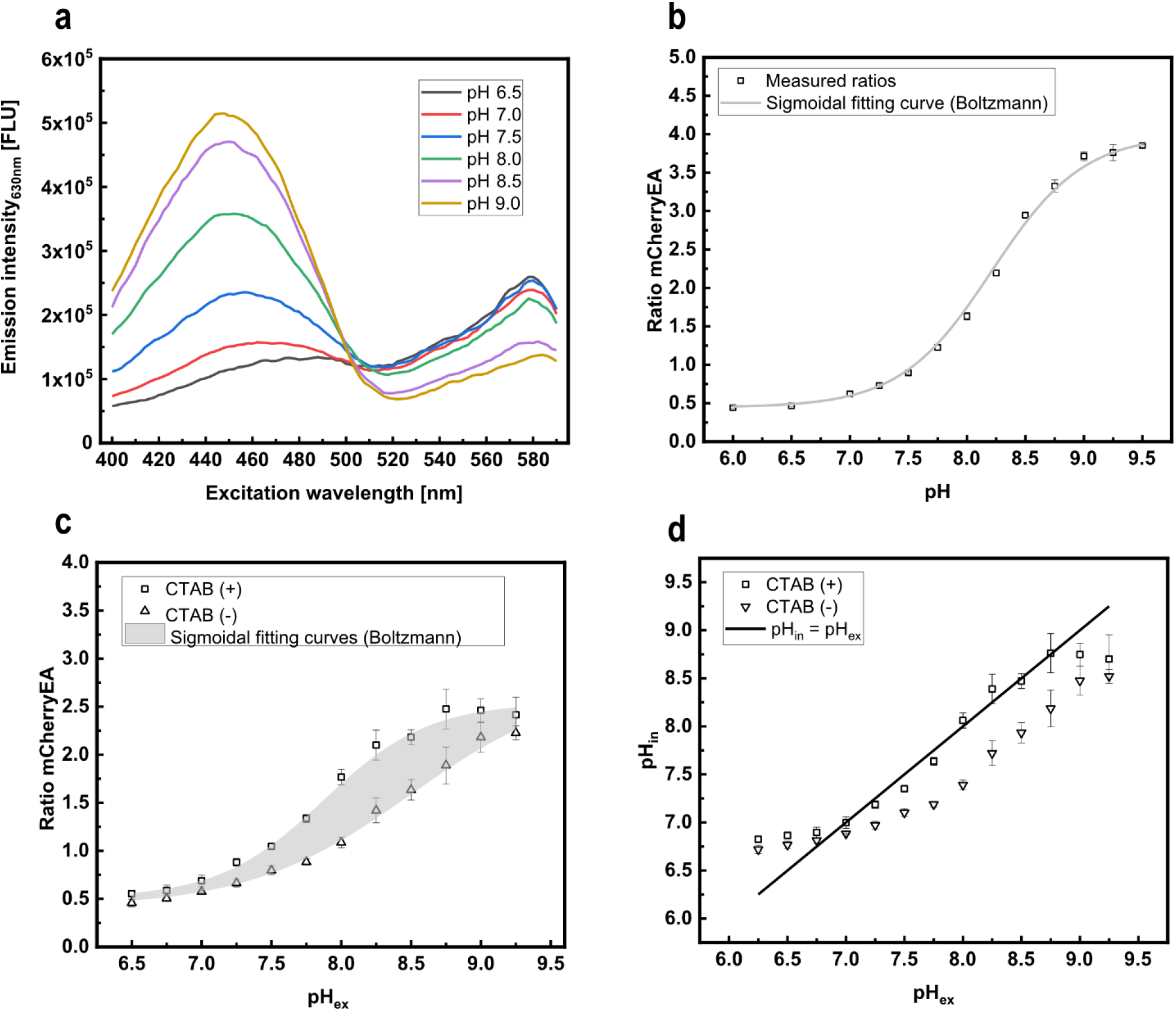
*In vitro* characterization of the pH biosensor mCherryEA using crude cell extracts of *E. coli* DH5α (pEKEx2_mCherryEA). The spectral biosensor response upon changes of the respective pH (a) and the corresponding calculated pH dependent ratios (b). mCherryEA biosensor ratio in *E. coli* DH5α (pEKEx2_mCherryEA) with and without the addition of CTAB (0.05% (w/v)) (c) and the calculated internal pH values of permeabilized cells compared to non- permeabilized cells (d). The biosensor protein was produced in *E. coli* DH5α (pEKEx2_mCherryEA). Cell extracts were prepared in 1 M PBS buffer with different pH values. For *in-situ* characterization, *E. coli* DH5α (pEKEx2_mCherryEA) was re-suspended in 1 M PBS buffer with different pH values and subsequently the cell suspension transferred to black 96- well plates. Fluorescence was measured before adding cetyltrimethylammonium bromide (CTAB) and after the addition of CTAB (incubation for 5 minutes in the dark prior repeating the fluorescence measurements). Ratio of the biosensor mCherryEA was calculated by dividing the emission intensity (630 nm) obtained with an excitation at 454 nm by an excitation of 580 nm. Error bars represent standard deviation calculated from at least three replicates. Curve fitting was conducted using a sigmoidal fit (Boltzmann) in Origin. Fluorescence measurements were conducted in a SpectraMax iD3 plate reader.

In order to test the pH biosensor mCherryEA for *in vivo* monitoring of internal pH values, we transformed *E. coli* MG 1655 as well as the triple mutant strain *E. coli* TK 2309, which lacks all three major potassium uptake systems (Trk- ,Kdp-, Kup-) with the sensor plasmid pXMJ19_*mCherryEA* resulting in sensor equipped strains *E coli* MG 1655 (pXMJ19_mCherryEA) (WT_S) and *E. coli* TK 2309 (pXMJ19_mCherryEA) (TK 2309_S). For *E. coli* TK 2309 a growth deficit was reported in presence of less than 5 mM of potassium in the growth media (Roe *et al.*, 2000). To verify this phenotype for the sensor carrying strain TK 2309_S, a growth experiment with WT_S as well as TK 2309_S was conducted in K_0.1_ and K_120_ minimal medium (Fig. 2a, b). Growth of the WT_S strain proceeded with a rate of 0.12 h^−1^ and 0.2 h^−1^ at 0.1 mM and 120 mM potassium, respectively. (Fig. 2a). At low potassium concentrations a growth deficit for TK 2309 was observed (Fig. 2b) (Roe *et al.*, 2000). However, this phenotype was abolished in presence of 120 mM potassium resulting in a growth rate of 0.22 h^−1^ (Fig. 2b).

**Figure 2:**
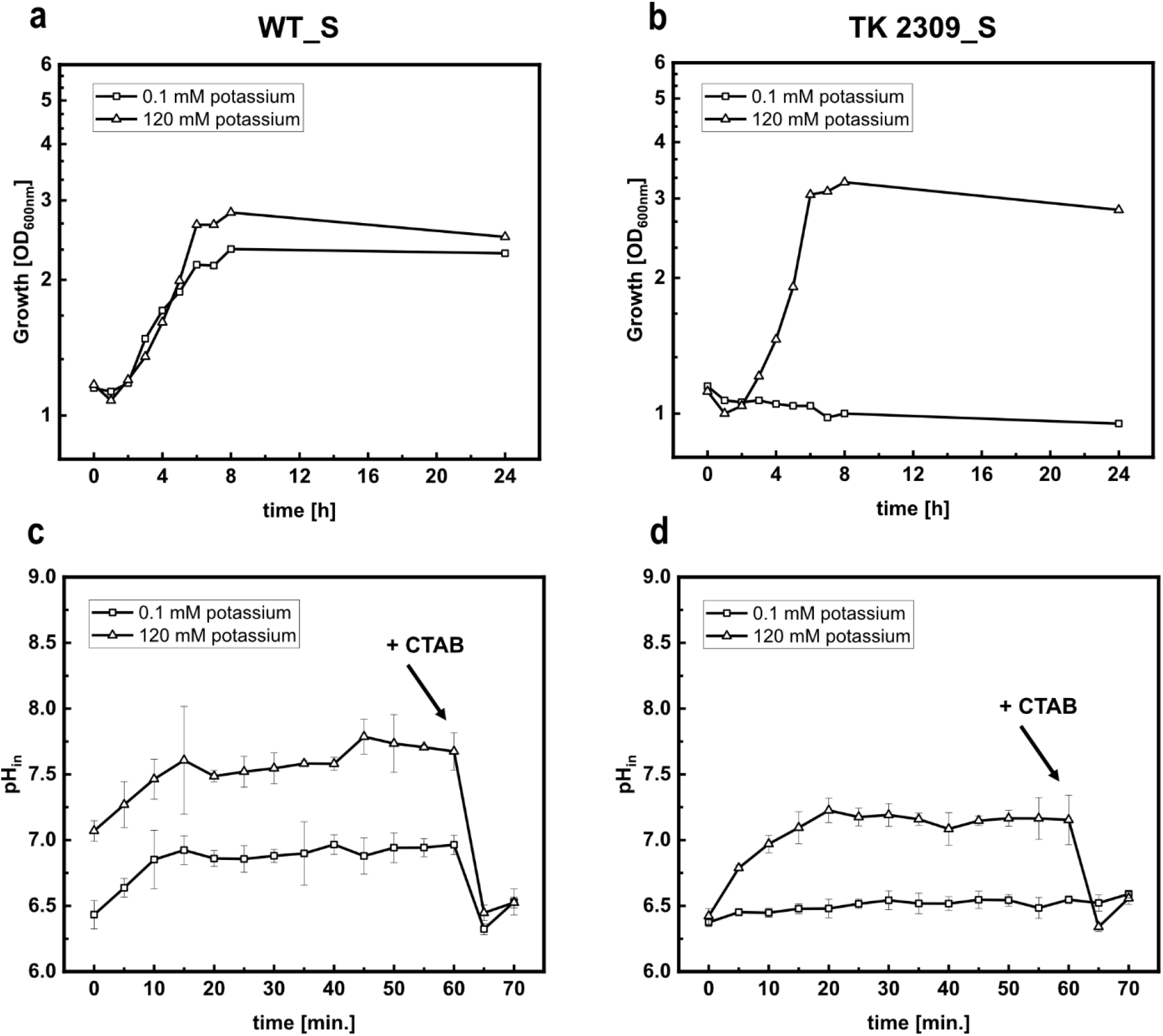
Growth (a, b) and *in-line* monitoring of the internal pH (c, d) using the biosensor mCherryEA in minimal medium supplemented with 0.1 mM potassium (K_0.1_) and 120 mM potassium (K_120_) for *E. coli* MG 1655 (pXMJ19_mCherryEA) (WT_S) (a, c) and *E. coli* TK 2309 (pXMJ19_mCherryEA) (TK 2309_S) (b, d). Growth experiment was conducted in 50 mL cultures (500 mL shaking flasks; 37°C, 150 rpm) using K_0.1_ and K_120_ minimal medium and 1 % glucose (w/v). Growth was monitored *via* the optical density at 600 nm. For *in-line* monitoring, biosensor strains were prepared as stated in the methods and materials part. Emission intensity at 630 nm was recorded upon an excitation at 454 nm and 580 nm. At the end of the experiment, cetyltrimethylammonium bromide (CTAB; final concentration 0.05% (w/v)) was added for sensor calibration purposes as it allows an equilibration of the internal and external environment.

For *E. coli* TK 2309 a strong shift of pH_in_ towards an acidic value of 6.3 has been described upon incubation at an external pH of 6 and low potassium concentrations (Roe *et al.*, 2000). This strong acidification of pH_in_ in *E. coli* TK 2309 does not occur in presence of 120 mM potassium and in *E. coli* strains with at least one functional potassium uptake system (Roe *et al.*, 2000). To re-investigate the effects of potassium on pH_in_ by using the biosensor mCherryEA, we adapted a recently described real-time pH-homeostasis experiment (Reyes-Fernández and Schuldiner, 2020). For this purpose, WT_S and TK 2309_S were pre-cultivated 24 hours in LK media followed by an cultivation step in K_30_ medium until the stationary phase was reached. Following that, the two strains were harvested, washed three times with 0.9 % NaCl and cultivated in K_0.1_ and K_120_ minimal medium for three hours. Finally, the cells were suspended in fresh K_0.1_ and K_120_ medium (pH of 6.0) containing 0.2 % (w/v) glucose and then immediately transferred as 180 μl aliquots into 96-well-plates. Cells were incubated at 37°C and fluorescence signals were recorded *in-line* for 60 minutes for determination of internal pH levels. As depicted in Fig. 2c, in presence of 0.1 mM potassium, the internal pH of WT_S increased from initially 6.5 to approximately 7.0 within 15 minutes of incubation. Upon addition of CTAB just before the experiment was ended the sensor signal for pH_in_ dropped from 7.0 to approximately 6.5, which corresponds to the lower detection limit of the pH biosensor mCherryEA. A similar time course for pH_in_ was observed for WT_S in presence of 120 mM potassium, for which the internal pH increased to approximately 7.7 after 15 minutes of incubation (Fig. 2c), before addition of CTAB caused the expected drop of the biosensor signal for pH_in_. The pH_in_ value of 7.7 measured for WT_S corresponds well to the pH_in_ values between 7.4 and 7.9 previously reported for *E. coli* WT strains when incubated under similar conditions (Roe *et al.*, 2000; Slonczewski *et al.*, 2009; Reyes-Fernández and Schuldiner, 2020). Investigation of pH_in_ *via* the sensor mCherryEA in the potassium uptake deficient strain TK2309_S revealed that the sensor signal remained constantly at the lower detection limit of 6.5 for incubation in K_0.1_ medium with pH 6.0 (Fig. 2d). As expected for this external pH, addition of CTAB at the end of the experiment did not have any impact on the sensor signal for pH_in_. In presence of 120 mM potassium, the initially recorded internal pH of 6.5 for TK2309_S increased within the first 20 minutes to around 7.3 ± 0.1 and remained stable at this level prior CTAB addition at the end of the experiment, which caused the expected drop of the sensor signal (Fig. 2d). The here detected pH_in_ values for TK2309_S below 6.5 in K_0.1_ medium and 7.3 for K_120_ medium correspond well with the internal pH values of 6.3 and 7.4 previously determined for this strain by the use of [7-^14^C]-benzoic acid (Roe *et al.*, 2000).

Taken together, these experiments illustrate well the functionality of the sensor mCherryEA for the analysis of pH_in_ in liquid cultures of *E. coli* but also revealed restrictions of this method, which are set by the lower and upper pH detection limits of 6.5 and 8.75 of the used sensor protein.

### 3.2 The biosensor protein mCherryEA can report the internal pH of *E. coli* colonies on agar plates

To test if the sensor can be used to directly assess internal pH levels in colonies on agar plates, colonies of *E. coli* MG 1655 (pXMJ19_mCherryEA) (WT_S) were spotted on rectangle plates containing SB agar supplemented with 1 mM IPTG. After the robot assisted spotting, the agar plates were incubated for 20 hours at 37 °C and the fluorescence of the colonies then detected *via* imaging using an Imaging system equipped for fluorescence analysis. The fluorescence detection was conducted using two different capsules for excitation (440 nm and 530 nm) and one filter (595 nm) to measure fluorescence emission. For colonies of WT_S, a mean fluorescence intensity of 2.22×10^4^ ± 552 FLU was measured for excitation at 440 nm and a higher fluorescence intensity of 3.04×10^4^ ± 717 FLU when excited at 530 nm (Fig. 3a). For the empty vector control strain *E. coli* (pXMJ19), colony fluorescence was more than four (0.48×10^4^ ± 90 FLU) and six (0.44×10^4^ ± 63 FLU) times lower upon excitation at 440 nm and 530 nm, respectively (Fig. 2S). These results show the proper expression of the biosensor protein mCherryEA in the WT_S colonies on SB agar plates supplemented with IPTG.

**Figure 3:**
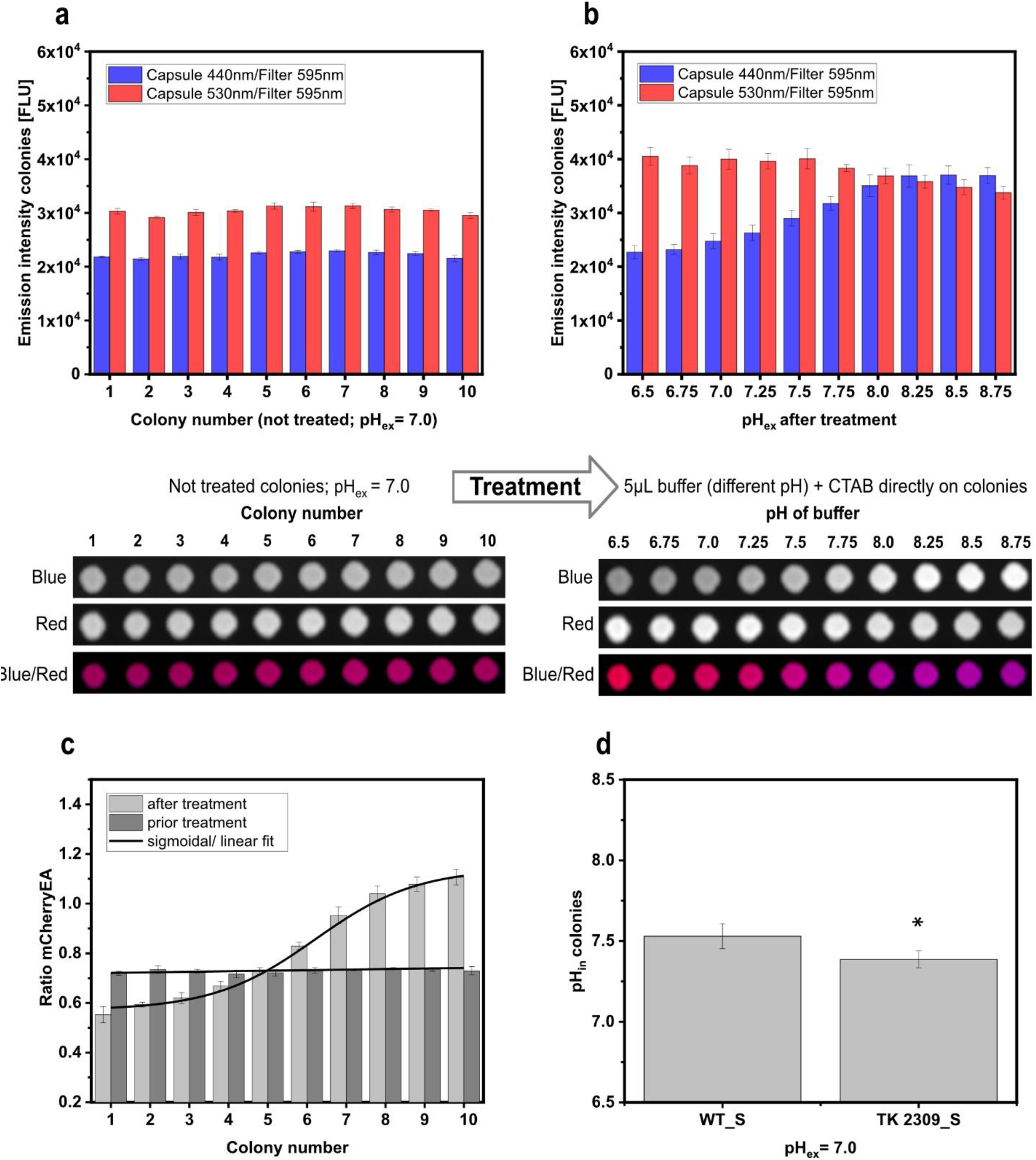
mCherryEA biosensor signals in *E. coli* MG 1655 (pXMJ19_mCherryEA) arrayed colonies on agar plate without any treatment (a) and after adding buffer with different pH values directly on the respective colonies (b). Calculated ratios prior and after treatment of the colonies (c) and determination of the internal pH values of *E. coli* MG 1655 (pXMJ19_mCherryEA) (WT_S) and *E. coli* TK 2309 (pXMJ19_mCherryEA) (TK 2309_S) (d). 1 M PBS buffer with different set pH values was supplemented with cetyltrimethylammonium bromide (CTAB; final concentration 0.05 % (w/v)). Internal pH values were analyzed with one-way-Anova followed by Tukey’s test (^n.s^ p > 0.05; * p ≤ 0.05). Error bars represent standard deviation from at least three replicates.

Genetically encoded FRPs have been shown to respond instantaneously to changes of the target internal parameter in liquid cultures as shown in this study for the FRP mCherryEA in *E. coli*. This property should allow to directly verify the biosensor functionality in colonies grown on agar plates. For this purpose, PBS buffer solutions with different set pH values containing 0.05 % CTAB were applied directly as 5 μl drops onto each of the colonies and then imaged again in the fluorescence imaging system. The changed fluorescence emission intensities after exposure of the colonies to the different buffer solutions revealed, that the biosensor mCherryEA in the now treated colonies responded in a ratiometric manner to the different pH values (Fig. 3b). In detail, the emission intensity at 595 nm for excitation at 440 nm increased when adding PBS buffer with higher pH values and the emission intensity for the excitation at 530 nm decreased at lower pH values (when compared to the initial intensities). By multiplexing the two fluorescence images derived for the treated colonies at different pH values, where excitation at 440 nm was assigned to the false color blue and excitation at 530 nm to the false color red, the effects of the exposure to low and high pH values could be visualized as a shift from red to blue colored colonies, respectively (Fig. 3b). The values for the fluorescence intensities obtained upon excitation at 440 nm and 530 nm determined for the colonies on the agar plate were used to calculate ratios for colonies before exposing them to the different set pH buffer solutions supplemented with CTAB and after their respective treatment. The ratio of the biosensor signal was shown to be in a narrow range (between 0.7 and 0.8) for all colonies before exposing them to the buffer solutions (Fig. 3c). Treatment of the colonies with the different pH-buffer solutions containing CTAB resulted in a two-fold ratio increase from 0.5-0.6 to 1.1-1.2 when adding buffer solutions with pH values of pH 6.5 or 8.75, respectively (Fig. 3c). Based on the biosensor ratios for untreated colonies, an internal pH of 7.53 ± 0.08 was determined for *E. coli* MG 1655 (pXMJ19_mCherryEA) (WT_S) (Fig. 3d). This is in accordance with an internal pH value of 7.43 ± 0.01 determined for liquid cultures of WT_S in SB-medium (liquid) (Fig. S3b). In addition, the internal pH of *E. coli* TK 2309 (pXMJ19_mCherryEA) (TK 2309_S) grown as colonies on agar plates was determined to be 7.39 ± 0.05 (Fig. 3d). An internal pH of 7.22 ± 0.02 was measured for TK 2309_S liquid cultures in SB-medium (Fig. S3b). By this means, that for both liquid cultures as well as the imaging method on agar plates, the internal pH of *E. coli* MG 1655 was significantly higher when compared to the TK 2309 mutant. Despite the differences with respect to their internal pH, no growth deficit was observed in liquid SB media for the TK2309_S strain when compared to the WT_S strain (Fig. S3a). Taken together, the results successfully revealed that:

i. the sensor protein mCherryEA is functional in colonies on agar plates and this method can be used to directly assess the pH_in_ from bacterial colonies on agar plates.
ii. imaging of the pH_in_ *via* the FRP (mCherryEA) signals from colonies on agar plates allows to distinguish colonies with impaired pH-homeostasis capabilities from those with a normal pH-homeostasis, under conditions which do not negatively affect growth patterns.

### 3.3 Fluorescent reporter protein-based screening of an *E. coli* transposon mutant library

To finally validate the concept of using a FRP- sensor to screen a strain library cultivated as colonies on agar plates, a Tn5 transposon mutant library of *E. coli* MG1655 was created and transformed with the plasmid pXMJ19_*mCherryEA*. Linker-PCR experiments revealed the expected heterogeneity of Tn5 insertion sites for isolates of the mutant library before and after transformation of the sensor plasmid (Fig. S4). Single colonies of the *E. coli* Tn5 mutant library carrying pXMJ19_*mCherryEA* were picked randomly and transferred to single wells filled with SB-medium in 96-well plates by the use of a QPix420 microbial colony picker. 96-well plates were cultivated over-night and then arrayed on SB-agar plates (pH 7.0, 1 mM IPTG) using a ROTOR HDA. After cultivation of the 384 clones arrayed on three SB-agar plates (128 each) at 37°C for 24 hours, the plates were imaged using the Vilber Fusion FX system. The average ratio of all colonies on the screening agar plates (Fig. 4a; Fig. S5a, b) was determined to be 0.50 ± 0.04. In Fig. 4a the sensor ratio distribution and respective fluorescence image of one screening plate is depicted. Sensor analysis revealed a colony possessing a reduced ratio of 0.44 (TP1) and another colony which ratio was drastically increased 0.78 (TP2). For all other colonies such a rational decision was not possible. However, one further colony located in the lower range of the colony ratios (TP 3; 0.48) and in the higher (TP 5; 0.59) were isolated for further analysis. Another interesting phenotype (Screening plate 1; Fig. S5a) was isolated due to its different morphological structure compared to the other colonies growing on the screening agar plate, even though the ratio of this mutant was just slightly increased with 0.55 (Fig. S5a). From screening plate 2 no mutant was selected, however, the fluorescence images as well as all analyzed ratios are provided in Fig. S5b.

**Figure 4:**
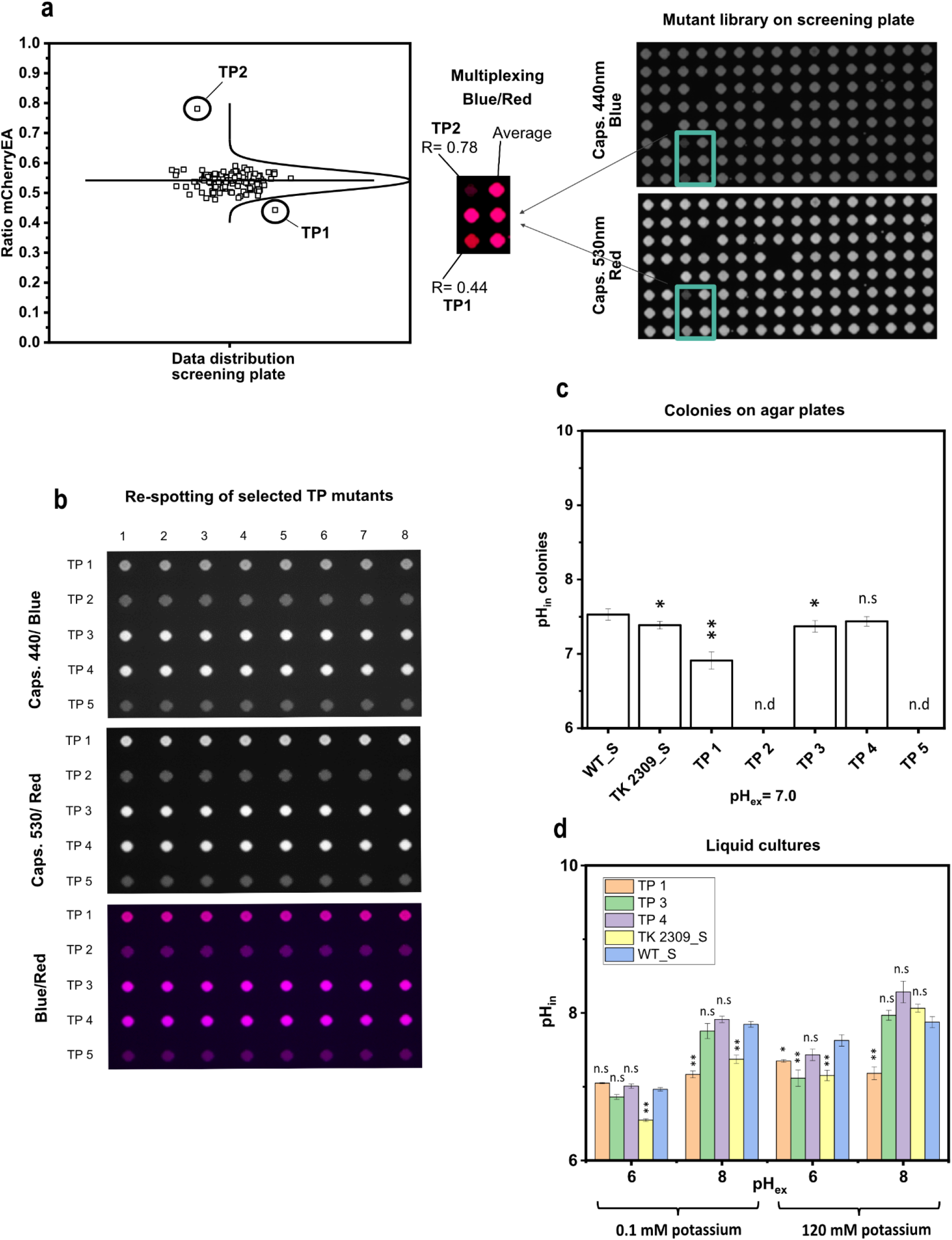
Fluorescence image of a screening plate with transposon derived mutants of *E coli* MG1655 equipped with the pH sensor plasmid pXMJ19_*mCherryEA* (a) and different selected transposon mutants re-spotted in eight replicates on SB-agar plates (b). The respective internal pH of eight replicates for the selected transposon mutants was determined and compared to *E. coli* MG 1655 (pXMJ19_*mCherryEA*) (WT_S) and *E. coli* TK 2309 (pXMJ19_*mCherryEA*) (TK 2309_S) (c). Internal pH levels of selected transposon mutants was verified and compared to WT_S and TK 2309_S strains in liquid media (minimal medium K_0.1_ and K_120_) at different external set pH values (d). Error bars represent standard deviation of at least three replicates. Statistical analysis was performed *via* One-Way-Anova followed by a Tukey’s test (^n.s^ p > 0.05; * p ≤ 0.05; ** p ≤ 0.01). Internal pH was not determined (n.d) for TP 2 and TP 5 mutants due to its weak biosensor expression.

The follow-up fluorescence imaging analysis of these five transposon mutants on SB-agar plates (pH 7.0; Fig. 4b) revealed indeed an acidification of the internal pH for *E. coli* TP 1 (pH_in_ of 6.91 ± 0.12) and *E. coli* TP 3 (pH_in_ 7.37 ± 0.08) when compared to the pH_in_ of 7.53 ± 0.08 measured for the reference strain *E. coli* MG 1655 (pXMJ19_*mCherryEA*) (Fig. 4c). No significant difference of pH_in_ in comparison to the reference strain was measured for colonies of *E. coli* TP 4 (pH_in_ 7.44 ± 0.06; Fig. 4c). In contrast to the sensor signals of *E. coli* mutants TP 1, TP 3 and TP 4, the detected biosensor signals for the mutants TP 2 and TP 5 were very low (Fig. 4b). This observation indicates that the biosensor gene was weakly expressed in the two mutants *E. coli* TP2 (TN-insertions at *cusF*) and *E. coli* TP5 (TN-insertion at *ypaD*), which does not allow reliable analysis of the ratiometric signals of the pH-sensor protein for these two strains (Fig. 4b, c).

For *E. coli* TP1 the Tn5 insertion was mapped to the *rssB* gene, which encodes the adaptor protein RssB, which is in the control of degradation of σ^S^ encoded by *rpoS* (Ruiz *et al.*, 2001; Pruteanu and Hengge-Aronis, 2002). For *E. coli* TP 3 the *trkH* gene for the major potassium uptake system TrkH (Schlosser *et al.*, 1995) was found to be disrupted by the transposon. For the Tn5-mutagenesis derived strain *E. coli* TP4, the gene *bcsA*, encoding the cellulose synthase BcsA (Srra *et al.*, 2013) was found to be disrupted. BcsA contributes to the synthesis of cellulose, a mayor structural composite required for biofilm formation (Serra *et al.*, 2013). Lack of BcsA could underlie the observed altered colony morphology of *E. coli* TP4 on the screening SB-agar plate (Fig. S5a). After re-spotting this strain, however, the morphologically changed structure of the colony was not reproducible but similar to the morphology of the other mutants. For BcsA, no involvement in pH-homeostasis has been reported. This is in agreement to the similar pH_in_-values measured for *E. coli* TP4 and the reference strain when cultivated on SB-agar plates (Fig. 4c).

To further validate the results, pH_in_ of the Tn5-mutants *E. coli* TP1, *E. coli* TP3 and *E. coli* TP4 was analyzed for liquid cultures in minimal medium at different pH_ex_ values and potassium concentrations after one hour of incubation (Fig. 4d). In presence of 120 mM potassium and a pH_ex_ of 6.0, pH_in_ values of 7.35 ± 0.02 and 7.15 ± 0.19 were measured for *E. coli* TP1 and *E. coli* TP3, respectively, which are significantly lower values when compared to a pH_in_ of 7.63 ± 0.08 measured for WT_S (Fig. 4d). In case of *E. coli* TP3, this value is almost identical to the pH_in_ value of 7.14 ± 0.15 of TK 2309_S (Fig. 4d). Upon exposure to a pH_ex_ of 8.0 and in presence of 120 mM potassium, the pH_in_ values of both *E. coli* TP3 and TK 2309_S mutant (pH_in_ 7.97 ± 0.07 and pH_in_ 8.06 ± 0.05, respectively) were not significantly different from the pH_in_ of 7.88 ± 0.08 measured for WT_S, whereas the pH_in_ of *E. coli* TP1 remained at 7.18 ± 0.09 (Fig. 4d). At low potassium concentrations (0.1 mM) and a pH_ex_ of 6.0 neutral to slightly acidic pH_in_ values were determined for all strains. In detail a pH_in_ of 7.05 ± 0.01 was measured for *E. coli* TP 1 and a pH_in_ of 6.96 ± 0.03 for WT_S, a pH_in_ of 6.86 ± 0.03 for *E. coli* TP 3, and the lowest pH_in_ of 6.55 ± 0.02 was measured in TK2309_S. To note, when setting an external pH of 8 and in presence of low potassium concentrations, still a neutral pH_in_ of 7.17 ± 0.1 was determined for *E. coli* TP 1. In contrast, under these conditions *E. coli* TP 3 and WT_S revealed more alkaline pH_in_ values of 7.75 ± 0.1 and 7.85 ± 0.04, respectively. A pH_in_ of 7.37 ± 0.06 was determined for *E. coli* TK 2309_S (Fig. 4d). As expected from the analyses on SB-agar plates, for all four tested cultivation conditions no significant differences of pH_in_ values of *E. coli* TP 4 when compared to these of the reference strain WT_S were observed (Fig. 4d).

These results show that the *E. coli* TP1 and *E. coli* TP3, identified *via* image analysis of colonies on SB-agar plates as candidates with altered pH-homeostasis properties, revealed Tn5 insertions in genes known to be involved in pH-homeostasis or pH-stress adaptation (Roe *et al.*, 2000; Battesti *et al.*, 2011). For the Tn5-mutant defective in *rssB* (TP1), it should be highlighted that this strain maintained a stable neutral pH_in_ between 7.0 and 7.2, independent of the here applied external conditions with respect to both pH_ex_ as well as potassium concentrations. This observed phenotype of *E. coli* TP1 is probably brought by constitutively high levels of the σ^S^ (RpoS), which in turn might lead to increased transcription of genes for the general stress response in *E. coli* (Battesti *et al.*, 2011; Gottesman, 2019). This can be explained by the functionality of rssB, as it mediates σ^S^ degradation by the ATP-dependent protease ClpXP in absence of any stress (Muffler *et al.*, 1996; Dorich *et al.*, 2019). Consequently, inactivation of the *rssB* gene leads to a constant “ON”-state of the general stress response because of the absence of the proteolytic degradation of RpoS. For *E. coli* TP3, which carries the Tn5 insertion within the *trkH* gene, it might irritate that the difference of pH_in_ (compared to the reference strain WT_S) was only detected to be significant in presence of high amounts of potassium. However, expression of genes for potassium systems is repressed in presence of high potassium concentrations in *E. coli* (Laermann *et al.*, 2013; Schramke *et al.*, 2016), which in turn causes alterations of pH_in_ in *trkH*-deficient strains also at potassium concentrations above 20 mM (Roe *et al.*, 2000).

Taken together, the use of a FRP sensor for internal pH measurements (mCherryEA) was successfully applied to identify mutants on agar plates with altered internal pH levels, which subsequently was verified in liquid cultures. This illustrates the potential and flexibility brought by FRP sensors as it can be applied for both real-time monitoring as well as screening purposes, two methods of great importance to understand intracellular processes mechanistically. The efficiency of a screening method is determined by its signal to noise ratio. The ratiometric characteristic of many FRPs, like the here applied mCherryEA, provide the advantage that the biosensor signal is independent of the absolute amount of sensor molecules and thus differences in expression strength within the library do not affect the sensor signal. To note, combinatorial sensor set-ups (TFBs and FRPs) could capture both actual intracellular values (metabolite concentration or physiological states) and more regulatory elements (activation or repression of genetic circuits). Gaining insights of both is key for a systematic and profound physiological characterization of engineered platform organisms for industrial Biotechnology. Thus, the here established method provides a first step towards implementing fast sensor proteins in a routinely manner at an early stage such as screening of mutant libraries for a better understanding of molecular mechanisms.

## 4 Conclusion

High-throughput arrays of bacterial strain libraries on agar plates have become a versatile tool for phenotypic analysis towards a comprehensive understanding of gene functions and interactions in microorganisms in various cultivation conditions (Côté *et al.*, 2016; Peters *et al.*, 2016; Galardini *et al.*, 2017). The use of genetically encoded sensors offer additional possibilities to investigate strain libraries for a single physiological parameter besides growth (Germond *et al.*, 2016; Sanford and Palmer, 2017; Sarnaik *et al.*, 2020) and the combination of transcription-factor-based biosensors (TFBs) with phenotypic arrays on agar plates enabled comprehensive, non-destructive, temporally resolved gene expression studies (French *et al.*, 2018). In contrast to this type of biosensor, fluorescent reporter proteins (FRPs) are commonly used in microorganisms for real-time monitoring of fast internal kinetics upon an environmental trigger (Bermejo *et al.*, 2011a; Martynov *et al.*, 2018; Depaoli *et al.*, 2019). On the one hand, their fast response is a challenge for their use in FACS- based screening approaches. On the other hand, this characteristic provides to measure the actual state of the cell rather than measuring physiological states or metabolite concentrations from the past. In this communication, the FRP- based genetically encoded pH-sensor mCherryEA was successfully applied to screen *E. coli* mutant colonies arrayed on agar plates *via* fluorescence imaging. By exchanging the filters of the imaging system towards the properties of other sensor proteins, the here developed technology can be applied for other metabolites and physiological states in colonies and thus become a versatile first step in comprehensive phenotypic analysis of genome-wide libraries of bacterial strains using FRP- based biosensors.

## Funding

This work was funded by the Novo Nordisk Fonden within the framework of the Fermentation- based Biomanufacturing Initiative (FBM) (FBM-grant: NNF17SA0031362).

## Availability of data and materials

All data generated and analyzed during this study are included in this article and its additional files. Raw datasets are available from the corresponding author on reasonable request.

## Conflict of Interest

The authors declare that the research was conducted in the absence of any commercial or financial relationships that could be construed as a potential conflict of interest.

## Acknowledgment

We would like to thank the Fermentation Core at DTU Bioengineering for excellent technical support and the members of the COST Action “Understanding and exploiting the impacts of low pH on microorganisms” (EuroMicropH) CA18113 for very helpful discussions.

## References

Battesti, A., Majdalani, N., and Gottesman, S. (2011) The RpoS-mediated general stress response in *Escherichia coli*. Annu Rev Microbiol 65: 189–213.

Bermejo, C., Ewald, J.C., Lanquar, V., Jones, A.M., and Frommer, W.B. (2011a) *In vivo* biochemistry: Quantifying ion and metabolite levels in individual cells or cultures of yeast. Biochem J 438: 1–10.

Bermejo, C., Haerizadeh, F., Takanaga, H., Chermak, D., and Frommer, W.B. (2011b) Optical sensors for measuring dynamic changes of cytosolic metabolite levels in yeast. Nat Protoc 6: 1806–1817.

Bertani, G. (2004) Lysogeny at mid-twentieth century: P1, P2, and other experimental systems. J Bacteriol 186: 595–600.

Blattner, F.R., Plunkett, G., Bloch, C.A., Perna, N.T., Burland, V., Riley, M., et al. (1997) The complete genome sequence of *Escherichia coli* K-12. Science 277: 1453–1462.

Botman, D., Heerden, J.H. van, and Teusink, B. (2020) An improved ATP FRET sensor for yeast shows heterogeneity during nutrient transitions. ACS sensors 5: 814–822.

Bushell, F., Herbert, J.M.J., Sannasiddappa, T.H., Warren, D., Keith Turner, A., Falciani, F., and Lund, P.A. (2021) Mapping the transcriptional and fitness landscapes of a pathogenic *E. coli* strain: The effects of organic acid stress under aerobic and anaerobic conditions. Genes 12: 53.

Cella, J.A., Eggenberger, D.N., Noel, D.R., Harriman, L.A., and Harwood, H.J. (1952) The relation of structure and critical concentration to the bactericidal activity of quaternary ammonium salts. J Am Chem Soc 74: 2061–2062.

Côté, J.P., French, S., Gehrke, S.S., MacNair, C.R., Mangat, C.S., Bharat, A., and Brown, E.D. (2016) The genome-wide interaction network of nutrient stress genes in *Escherichia coli*. MBio 7: e01714–16.

Crauwels, P., Schäfer, L., Weixler, D., Bar, N.S., Diep, D.B., Riedel, C.U., and Seibold, G.M. (2018) Intracellular pHluorin as sensor for easy assessment of bacteriocin-induced membrane-damage in *Listeria monocytogenes*. Front Microbiol 9: 1–10.

Deng, H., Li, J., Zhou, Y., Xia, Y., Chen, C., Zhou, Z., et al. (2021) Genetic engineering of circularly permuted yellow fluorescent protein reveals intracellular acidification in response to nitric oxide stimuli. Redox Biol 41: 101943.

Depaoli, M.R., Bischof, H., Eroglu, E., Burgstaller, S., Ramadani-Muja, J., Rauter, T., et al. (2019) Live cell imaging of signaling and metabolic activities. Pharmacol Ther 202: 98–119.

Dorich, V., Brugger, C., Tripathi, A., Hoskins, J.R., Tong, S., Suhanovsky, M.M., et al. (2019) Structural basis for inhibition of a response regulator of σS stability by a ClpXP antiadaptor. Genes Dev 33: 718–732.

Eggeling, L., Bott, M., and Marienhagen, J. (2015) Novel screening methods-biosensors. Curr Opin Biotechnol 35: 30–36.

Eikmanns, B.J., Kleinertz, E., Liehl, W., and Sahm, H. (1991) A family of *Corynebacterium glutamicum*/*Escherichia coli* expression and promoter probing. Plasmid 102: 93–98.

Epstein, W. (1986) Osmoregulation by potassium transport in *Escherichia coli*. FEMS Microbiol Lett 39: 73–78.

Epstein, W., and Kim, B.S. (1971) Potassium transport loci in *Escherichia coli* K-12. J Bacteriol 108: 639–644.

French, S., Coutts, B.E., and Brown, E.D. (2018) Open-source high-throughput phenomics of bacterial promoter-reporter strains. Cell Syst 7: 339–346.

Galardini, M., Koumoutsi, A., Herrera-Dominguez, L., Varela, J.A.C., Telzerow, A., Wagih, O., et al. (2017) Phenotype inference in an *Escherichia coli* strain panel. Elife 6: e31035.

Germond, A., Fujita, H., Ichimura, T., and Watanabe, T.M. (2016) Design and development of genetically encoded fluorescent sensors to monitor intracellular chemical and physical parameters. Biophys Rev 8: 121–138.

Goldbeck, O., Eck, A.W., and Seibold, G.M. (2018) Real time monitoring of NADPH concentrations in *Corynebacterium glutamicum* and *Escherichia coli* via the genetically encoded sensor mBFP. Front Microbiol 9: 1–10.

Gottesman, S. (2019) Trouble is coming: Signaling pathways that regulate general stress responses in bacteria. J Biol Chem 294: 11685–11700.

Guan, N., and Liu, L. (2020) Microbial response to acid stress: mechanisms and applications. Appl Microbiol Biotechnol 104: 51–65.

Guerrero-Castro, J., Lozano, L., and Sohlenkamp, C. (2018) Dissecting the acid stress response of *Rhizobium tropici* CIAT 899. Front Microbiol 9: 846.

Han, J., and Burgess, K. (2010) Fluorescent indicators for intracellular pH. Chem Rev 110: 2709–2728.

Hartmann, F.S.F., Clermont, L., Tung, Q.N., Antelmann, H., and Seibold, G.M. (2020) The industrial organism *Corynebacterium glutamicum* requires mycothiol as antioxidant to resist against oxidative stress in bioreactor cultivations. Antioxidants 9: 1–13.

Hayes, E.T., Wilks, J.C., Sanfilippo, P., Yohannes, E., Tate, D.P., Jones, B.D., et al. (2006) Oxygen limitation modulates pH regulation of catabolism and hydrogenases, multidrug transporters and envelope composition in *Escherichia coli* K-12. BMC Microbiol 6: 1–18.

Heins, A.L., Reyelt, J., Schmidt, M., Kranz, H., and Weuster-Botz, D. (2020) Development and characterization of *Escherichia coli* triple reporter strains for investigation of population heterogeneity in bioprocesses. Microb Cell Fact 19: 1–20.

Ito, M., Morino, M., and Krulwich, T.A. (2017) Mrp antiporters have important roles in diverse bacteria and archaea. Front Microbiol 8: 2235.

Jakoby, M., Carole-Estelle, Ngouoto-Nkili, and Burkovski, A. (1999) Construction and application of new *Corynebacterium glutamicum* vectors. Biotechnol Tech 13: 437–441.

Kashket, E.R. (1985) The proton motive force in bacteria: A critical assessment of methods. Annu Rev Microbiol 39: 219–242.

Koch, M., Pandi, A., Borkowski, O., Cardoso Batista, A., and Faulon, J.L. (2019) Custom-made transcriptional biosensors for metabolic engineering. Curr Opin Biotechnol 59: 78–84.

Krulwich, T.A., Sachs, G., and Padan, E. (2011) Molecular aspects of bacterial pH sensing and homeostasis. Nat Rev Microbiol 9: 330–343.

Laermann, V., Ćudić, E., Kipschull, K., Zimmann, P., and Altendorf, K. (2013) The sensor kinase KdpD of *Escherichia coli* senses external K+. Mol Microbiol 88: 1194–1204.

Lund, P., Tramonti, A., and Biase, D. De (2014) Coping with low pH: Molecular strategies in neutralophilic bacteria. FEMS Microbiol Rev 38: 1091–1125.

Lund, P.A., Biase, D. De, Liran, O., Scheler, O., Mira, N.P., Cetecioglu, Z., et al. (2020) Understanding how microorganisms respond to acid pH is central to their control and successful exploitation. Front Microbiol 11: 2233.

Maglica, Ž., Özdemir, E., and McKinney, J.D. (2015) Single-cell tracking reveals antibiotic-induced changes in mycobacterial energy metabolism. MBio 6: e02236–14.

Martinez, K.A., Kitko, R.D., Mershon, J.P., Adcox, H.E., Malek, K.A., Berkmen, M.B., and Slonczewski, J.L. (2012) Cytoplasmic pH response to acid stress in individual cells of *Escherichia coli* and *Bacillus subtilis* observed by fluorescence ratio imaging microscopy. Appl Environ Microbiol 78: 3706–14.

Martynov, V.I., Pakhomov, A.A., Deyev, I.E., and Petrenko, A.G. (2018) Genetically encoded fluorescent indicators for live cell pH imaging. Biochim Biophys Acta - Gen Subj 1862: 2924–2939.

Maurer, L.M., Yohannes, E., Bondurant, S.S., Radmacher, M., and Slonczewski, J.L. (2005) pH regulates genes for flagellar motility, catabolism, and oxidative stress in *Escherichia coli* K-12. J Bacteriol 187: 304–319.

Miesenböck, G., Angelis, D.A. De, and Rothman, J.E. (1998) Visualizing secretion and synaptic transmission with pH-sensitive green fluorescent proteins. Nature 394: 192–195.

Mira, N.P., Palma, M., Guerreiro, J.F., and Sá-Correia, I. (2010) Genome-wide identification of *Saccharomyces cerevisiae* genes required for tolerance to acetic acid. Microb Cell Fact 9: 1–13.

Monteiro, F., Hubmann, G., Takhaveev, V., Vedelaar, S.R., Norder, J., Hekelaar, J., et al. (2019) Measuring glycolytic flux in single yeast cells with an orthogonal synthetic biosensor. Mol Syst Biol 15: e9071.

Muffler, A., Fischer, D., Altuvia, S., Storz, G., and Hengge-Aronis, R. (1996) The response regulator RssB controls stability of the σS subunit of RNA polymerase in *Escherichia coli*. EMBO J 15: 1333–1339.

Palud, A., Scornec, H., Cavin, J.F., and Licandro, H. (2018) New genes involved in mild stress response identified by transposon mutagenesis in *Lactobacillus paracasei*. Front Microbiol 9: 535.

Peters, J.M., Colavin, A., Shi, H., Czarny, T.L., Larson, M.H., Wong, S., et al. (2016) A comprehensive CRISPR-based functional analysis of essential genes in bacteria. Cell 165: 1493–1506.

Piatkevich, K.D., Malashkevich, V.N., Almo, S.C., and Verkhusha, V. V. (2010) Engineering ESPT pathways based on structural analysis of LSSmKate red fluorescent proteins with large stokes shift. J Am Chem Soc 132: 10762–10770.

Pruteanu, M., and Hengge-Aronis, R. (2002) The cellular level of the recognition factor RssB is rate-limiting for σS proteolysis: Implications for RssB regulation and signal transduction in σS turnover in *Escherichia coli*. Mol Microbiol 45: 1701–1713.

Rajendran, M., Claywell, B., Haynes, E.P., Scales, U., Henning, C.K., and Tantama, M. (2018) Imaging pH dynamics simultaneously in two cellular compartments using a ratiometric pH-sensitive mutant of mCherry. ACS Omega 3: 9476–9486.

Reva, O.N., Weinel, C., Weinel, M., Böhm, K., Stjepandic, D., Hoheisel, J.D., and Tümmler, B. (2006) Functional genomics of stress response in *Pseudomonas putida* KT2440. J Bacteriol 188: 4079–4092.

Reyes-Fernández, E.Z., and Schuldiner, S. (2020) Acidification of cytoplasm in *Escherichia coli* provides a strategy to cope with stress and facilitates development of antibiotic resistance. Sci Rep 10: 1–13.

Roe, A.J., McLaggan, D., O’Byrne, C.P., and Booth, I.R. (2000) Rapid inactivation of the *Escherichia coli* Kdp K+ uptake system by high potassium concentrations. Mol Microbiol 35: 1235–1243.

Ruiz, N., Peterson, C.N., and Silhavy, T.J. (2001) RpoS-dependent transcriptional control of sprE: Regulatory feedback loop. J Bacteriol 183: 5974–5981.

Sambrook, J., Fritsch, E.F., and Maniatis, T. (1989) Molecular cloning: a laboratory manual. No. Ed. 2., Cold spring harbor laboratory press, .

Sanford, L., and Palmer, A. (2017) Recent advances in development of genetically encoded fluorescent sensors. Methods Enzymol 589: 1–49.

Sarnaik, A., Liu, A., Nielsen, D., and Varman, A.M. (2020) High-throughput screening for efficient microbial biotechnology. Curr Opin Biotechnol 64: 141–150.

Schallmey, M., Frunzke, J., Eggeling, L., and Marienhagen, J. (2014) Looking for the pick of the bunch: high-throughput screening of producing microorganisms with biosensors. Curr Opin Biotechnol 26: 148–154.

Schlosser, A., Meldorf, M., Stumpe, S., Bakker, E.P., and Epstein, W. (1995) TrkH and its homolog, TrkG, determine the specificity and kinetics of cation transport by the Trk system of *Escherichia coli*. J Bacteriol 177: 1908–1910.

Schramke, H., Tostevin, F., Heermann, R., Gerland, U., and Jung, K. (2016) A dual-sensing receptor confers robust cellular homeostasis. Cell Rep 16: 213–221.

Serra, D.O., Richter, A.M., and Hengge, R. (2013) Cellulose as an architectural element in spatially structured *Escherichia coli* biofilms. J Bacteriol 195: 5540–5554.

Shin, J., Zhang, S., Der, B.S., Nielsen, A.A.K., and Voigt, C.A. (2020) Programming *Escherichia coli* to function as a digital display. Mol Syst Biol 16: 1–12.

Slonczewski, J.L., Fujisawa, M., Dopson, M., and Krulwich, T.A. (2009) Cytoplasmic pH measurement and homeostasis in bacteria and archaea. Adv Microb Physiol 55: 1–317.

Studier, F.W., and Moffatt, B.A. (1986) Use of bacteriophage T7 RNA polymerase to direct selective high-level expression of cloned genes. J Mol Biol 189: 113–130.

Tantama, M., Hung, Y.P., and Yellen, G. (2011) Imaging intracellular pH in live cells with a genetically encoded red fluorescent protein sensor. J Am Chem Soc 133: 10034–10037.

